# Synergistic coevolution accelerates genome evolution

**DOI:** 10.1101/2021.05.19.444833

**Authors:** Daniel Preussger, Alexander Herbig, Christian Kost

**Affiliations:** Experimental Ecology and Evolution Research Group, Department of Bioorganic Chemistry, Max Planck Institute for Chemical Ecology, 07745 Jena, Germany; Department of Ecology, School of Biology/Chemistry, Osnabrück University, Osnabrück, 49076, Germany; Institute of Molecular Pathogenesis, ‘Friedrich-Loeffler-Institut’ (Federal Institute for Animal Health), Naumburger Str. 96a, 07743 Jena, Germany; Department of Archaeogenetics, Max Planck Institute for the Science of Human History, 07745 Jena, Germany; Department of Archaeogenetics, Max Planck Institute for Evolutionary Anthropology, 04103 Leipzig, Germany

## Abstract

Ecological interactions are key drivers of evolutionary change. Although it is well-documented that antagonistic coevolution can accelerate molecular evolution, the evolutionary consequences of synergistic coevolution remain poorly understood. Here we show experimentally that also synergistic coevolution can speed up the rate of molecular evolution. Pairs of auxotrophic genotypes of the bacterium *Escherichia coli*, whose growth depended on a reciprocal exchange of amino acids, were experimentally coevolved, and compared to two control groups of independently growing cells. Coevolution drove the rapid emergence of a strong metabolic cooperation that correlated with a significantly increased number of mutations in coevolved auxotrophs as compared to monoculture controls. These results demonstrate that synergistic coevolution can cause rapid evolution that in the long run may drive diversification of mutualistically interacting species.

**One-Sentence Summary:** Synergistic coevolution among obligate mutualists increases the rate of molecular evolution relative to independent types.

## Main Text

Identifying the factors that drive molecular change in the genomes of organisms is one of the most central goals of evolutionary biology (*1, 2*). Past research suggests that biotic interactions are fundamental for determining the evolution of organisms in nature (*3, 4*). Individuals displaying phenotypes that are advantageous in the context of a certain ecological interaction will be favoured by natural selection, while others lacking the corresponding alleles will be selected against (*5*). In cases where selection pressures are reciprocal, ecological interactions even set off coevolutionary dynamics of adaptation and counter-adaptation that can lead to perpetually escalating arms-races among groups of interacting individuals (*6*). As a consequence, ecological interactions do not only strongly affect the Darwinian fitness of the organisms involved, but also drive the molecular evolution of their genomes.

This causal link is well-established for antagonistic interactions, such as between parasites and their hosts (*7, 8*) or predators and their prey (*9*). For both of these cases, the so-called Red Queen hypothesis predicts adaptations that increase the fitness of one of the two interacting partners should simultaneously decrease the fitness of its corresponding counterpart (*10*–*13*). From this process results a strong evolutionary pressure that forces both sides to constantly evolve new adaptive phenotypes in order to survive and reproduce. Both comparative (*14, 15*) and experimental studies (*7, 16, 17*) corroborate that antagonistic coevolution increases the rate of molecular evolution, particularly affecting specific loci that are driving the interaction (*14, 15, 18*).

In contrast, little is known on how mutualistic coevolution affects the genomes of both interacting partners. Even though it is clear that engaging in mutualistic interactions can drastically affect the fitness of the organisms involved (*19*), the question how mutualistic cooperation affects the speed of molecular evolution, remains unanswered.

Two possibilities are conceivable. First and analogous to antagonistic coevolution, also in synergistic interactions individuals may be faced with a constantly evolving partner and, thus, experience the need to continuously respond to the accruing changes. In this case, synergistic coevolution would speed up molecular evolution. Alternatively, interacting partners may converge to a point of evolutionary stasis (*20, 21*), at which a unilateral change in one of the two parties is going to negatively affect the interaction as a whole. Under these conditions, stabilizing selection should maintain phenotypes of mutualists over extended periods of time, which, as a consequence, should decelerate the rate of molecular evolution.

Comparative studies analysing this issue by focusing on one side of a mutualistic association found evidence for an increased rate of molecular evolution in mutualistic species as compared to closely related non-mutualists existing outside the interaction (*22-24*). However, a direct empirical test of how synergistic coevolution affects the rate of molecular evolution in two interacting partners is lacking.

Here, we address this issue using experimental evolution of replicated populations of the bacterium *Escherichia coli*. Serially propagating two auxotrophic genotypes of this species (i.e. Δ*tyrA* and Δ*trpB*) resulted in the rapid evolution of a costly metabolic cooperation (*25*). Only 150 generations of coevolution transformed the initial consortia, in which both strains could only grow when they reciprocally exchanged essential amino acids, into a truly cooperative interaction, in which both parties started to actively invest costly resources to benefit their corresponding counterpart. This evolutionary transition was enabled by the formation of multicellular clusters among auxotrophic genotypes, which not only facilitated an exchange of amino acids between interaction partners (*26, 27*), but also immediately rewarded an increased cooperative investment of either party by positive fitness feedbacks (*25*). Importantly, these patterns were neither observed in auxotrophs that were cultivated individually in the presence of the required amino acid, nor in populations of metabolically autonomous (i.e. prototrophic) wild type cells, which were both propagated the same way as cocultured auxotrophs. This experimental design afforded the separation of effects resulting from the adaptation to the abiotic environment from those stemming from an adaptation to the cocultured partner.

After the evolution experiment, whole-population samples and individually isolated strains of all three experimental groups (i.e. (i) monocultures of prototrophic wild type (WT), (ii) monocultures of auxotrophic genotypes (M), and (iii) cocultures of auxotrophic genotypes (C)) were subjected to high-coverage, second-generation whole-genome re-sequencing to determine the number and identity of mutations that arose in the course of the evolution experiment. For this, derived genomes of all three treatment-groups were compared to the genomes of their corresponding evolutionary ancestor.

In this analysis, coevolved auxotrophs showed significantly increased numbers of mutations as compared to the two control groups (Fig. 1).

**Fig. 1.**
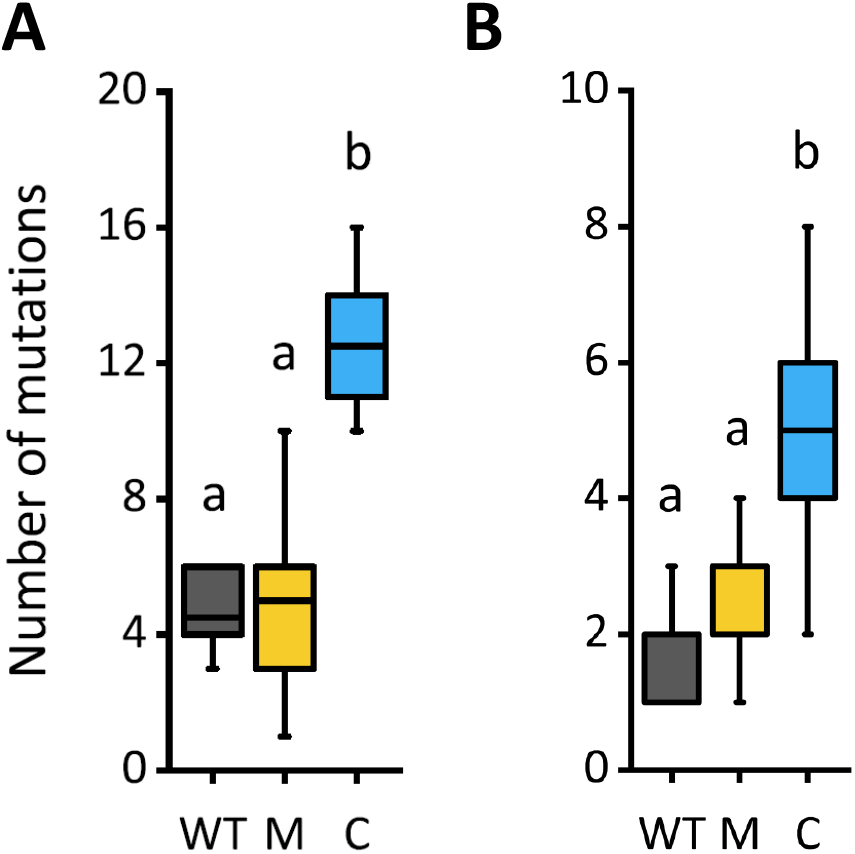
Coevolved auxotrophs accumulated more mutations than control groups. Absolute numbers of mutations of derived populations of wild type (WT), auxotrophic monocultures (M), and cocultured auxotrophs (C). Comparison of experimental groups was performed on the level of (**A**) whole population samples (Bonferroni posthoc test: P < 0.05; WT: n=6, M: n=11, C: n=6) and (**B**) samples of isolated clones (Bonferroni posthoc test: P < 0.05; WT: n=10, M: n=20, C: n=31). Different letters indicate significant differences between groups.

This was true on both the strain- and the population-level: median numbers (± 95% confidence interval) of mutations detected in coculture samples were more than twice as high as the ones detected in both control groups (i.e. population-level: C: 12.5 ± 1.6, WT: 4.5 ± 1.8, M: 5.0 ± 1.4; strain-level: C: 5.0 ± 0.5, WT: 2.0 ± 0.5, M: 2.0 ± 0.9). This observation is consistent with the interpretation that synergistic coevolution in auxotrophic cocultures accelerated the rate of molecular evolution relative to control populations that were able to grow independently. Additionally, statistically analysing clonal samples showed that the obligate interaction had a significant effect on the accumulated number of mutations (univariate linear model: P < 0.05, observed power: 0.85, WT: n=10, M: n=20, C: n=31).

To analyse the degree of divergence among derived genomes, gene-level distance trees were constructed by comparing the spectrum of mutations detected within samples. These trees revealed on both the level of isolated strains (Fig. 2) and entire populations (Fig. S1) that coevolved auxotrophs showed much higher levels of genetic divergence than the two control groups. This pattern was further corroborated by comparing pairwise distances based on the branch lengths among groups of isolated strains, which were significantly increased in coculture-derived samples relative to the ones of control groups (Dunnett T3 posthoc test: P < 0.05). Strikingly, auxotrophic cocultures formed independent branches in the tree, which were clearly distinct from the ones of control groups (Figs. 2 and S1). Depending on their auxotrophy-causing mutation, the two different cocultured genotypes (i.e. Δ*tyrA* and Δ*trpB*) even segregated into completely disparate branches, which suggests specialisation in the context of the mutually beneficial interaction. In contrast, this pattern was not observed in control groups. Here, derived populations of the prototrophic wild type and monocultures of the two auxotrophs formed intermixed branches without a clear genotypic separation. These findings strongly suggest (i) general differences in the spectrum of mutations between auxotrophic cocultures and control groups, (ii) highly divergent evolutionary trajectories between the two cooperating partners within a mutualistic consortium, and (iii) increased levels of parallel evolution in samples from cocultures relative to control groups.

**Fig. 2.**
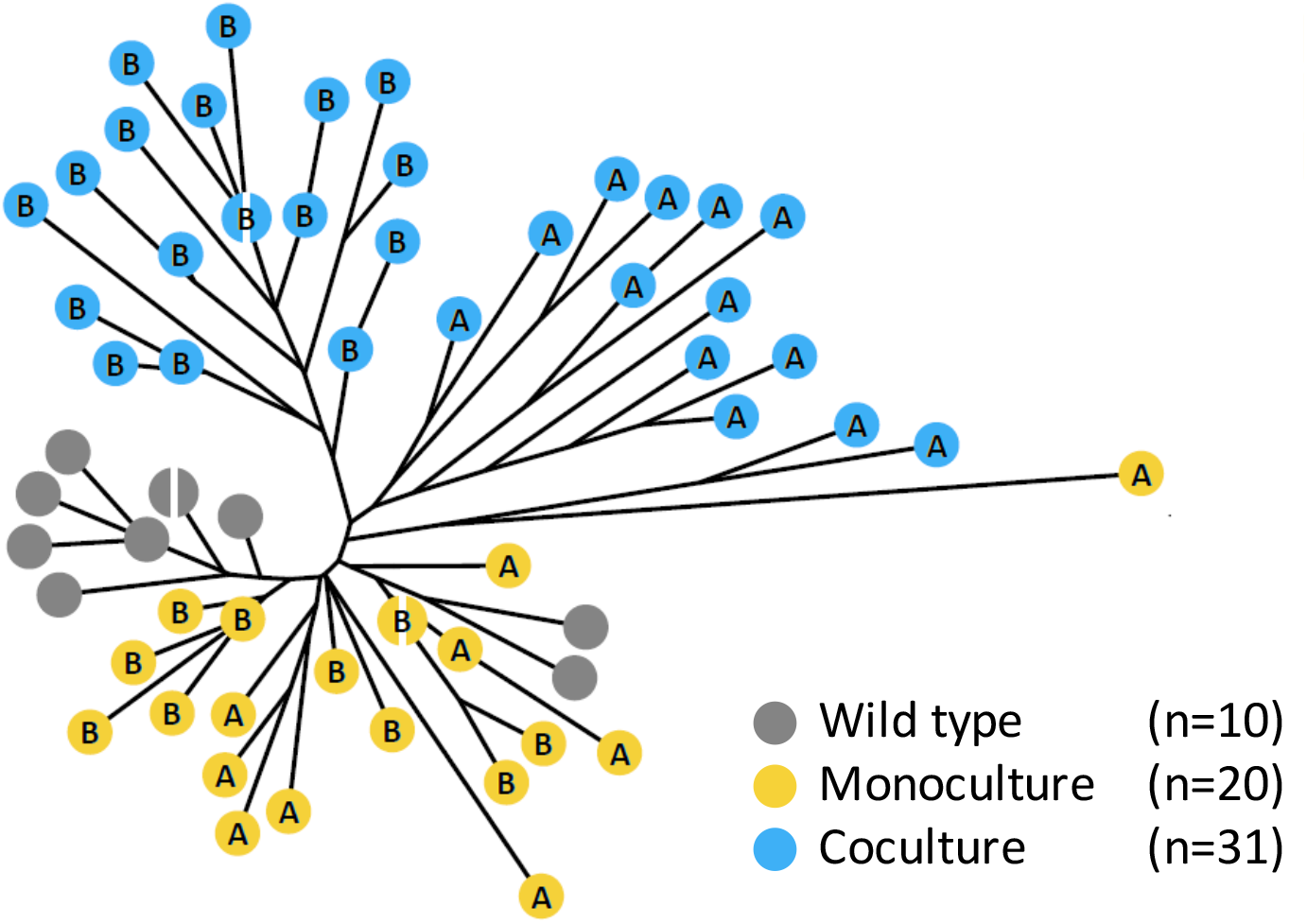
Coevolved auxotrophs showed an increased partner-specific divergence relative to control groups. Distance tree of isolated strains (A = tyrosine auxotroph, B = tryptophan auxotroph) that is based on the neighbour-joining analysis of a gene-level genotypic matrix (i.e. genes carrying a mutation were treated as alleles different to the wild type background). Split nodes indicate two samples mapped to the same position.

To further investigate whether the three treatment groups indeed differed in their degree of parallel evolution, the Jaccard index was calculated on the level of mutated genes for clonal samples for all possible comparisons within and between experimental groups (Fig. 3). Because growth of the two cocultured auxotrophic genotypes was limited by the availability of the required amino acid, the main route to increase in fitness was a mutation that enhanced growth of the other auxotrophic genotypes that were part of the same cluster. In this way, cooperative mutants automatically received more of the required amino acid in return (i.e. a positive feedback loop (*25*)). Hence, due to strong selection pressures that likely acted on cocultures of auxotrophs, analysed genomes should display a high degree of parallel evolution between replicate populations. In contrast, monocultures of auxotrophs and prototrophic wild type cells grew autonomously. Thus, the fitness effect of any mutation that improved a strain’s growth and survival in the current abiotic environment had to be strong enough in order to persist against and potentially outcompete other genotypes. Given the comparably short duration of the selection experiment (i.e. ∼ 153 generations), a much lower degree of parallel evolution was expected for these samples. Testing this hypothesis indeed revealed that coevolved auxotrophs showed a significantly increased degree of parallel evolution as compared to the two control groups (Fig. 3). This pattern emerged on a population-level (Fig. 3A) as well as when genotypes of a similar auxotrophy were compared between monoculture and coculture conditions (Fig. 3B). The generally increased degree of parallel evolution in cocultured auxotrophs indicates that these populations were likely subject to increased selection pressures.

**Fig. 3.**
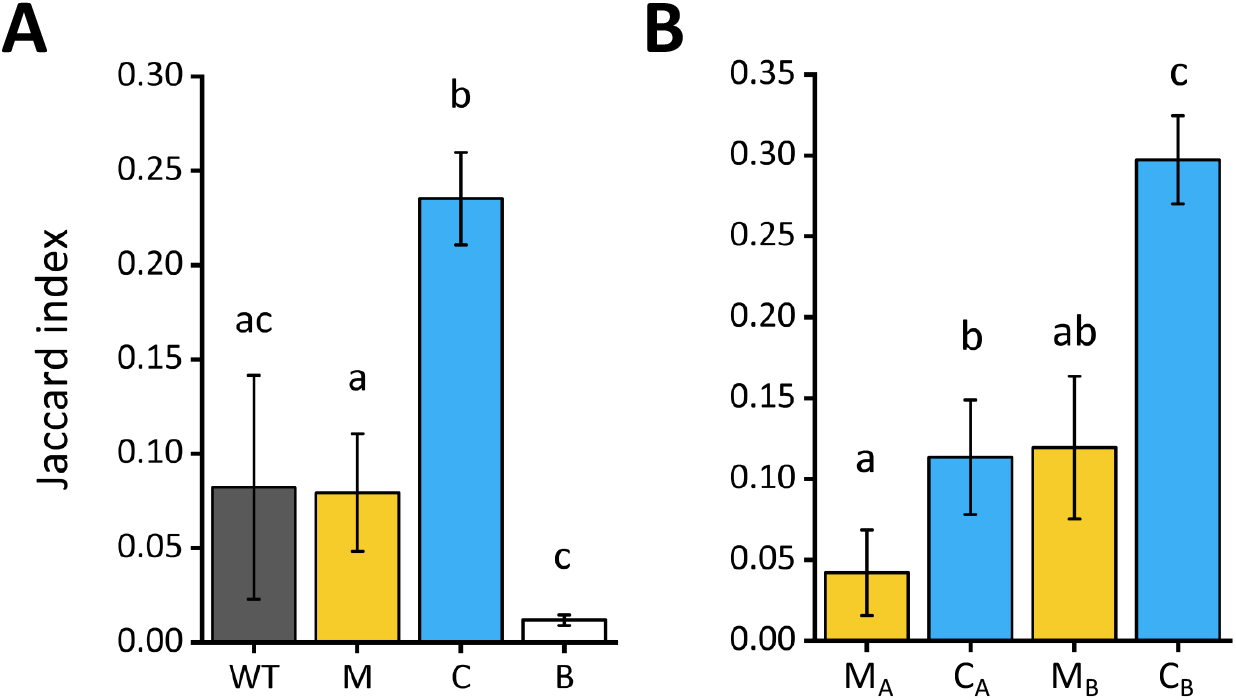
Coevolved auxotrophs show an increased degree of parallel evolution relative to control groups. Degree of parallel evolution within and between experimental groups based on mutated genes in sequenced isolates is given as mean Jaccard indices (± 95% confidence interval). (**A**) Comparison between populations of wild type (WT, n=45), monocultures of auxotrophs (M, n=110), cocultures of auxotrophs (C, n=231), and all experimental groups (B, n=1567). Different letters indicate significant differences between group (Dunnett T3 posthoc test: P < 0.05). (**B**) Comparison between isolates of monocultures (M_A, n=11; M_B, n=11) and cocultures (C_A: n= 13, C_B: n=18) on the level of the two individual auxotrophies (A: tyrosine auxotrophy (Δ*tyrA*); B: tryptophan auxotrophy (Δ*trpB*)). Different letters indicate significant differences between groups (Dunnett T3 posthoc test: *P* < 0.01).

In contrast, the degree of parallel evolution between groups of samples was generally low (Fig. 3) with very few mutational targets being shared between groups (Fig. S2). Of all 117 mutated genes identified on a population-level, only one gene was found to be mutated in all three experimental groups (Fig. S2A). Also, comparing the mutational spectra of auxotrophs evolved under monoculture and coculture conditions revealed that very few mutated genes were shared between groups (Δ*tyrA*: 3%, Δ*trpB*: 0%, Fig. S2B). Moreover, analysing the mutational data confirmed the previously observed high degree of divergence between cocultured auxotrophs (Fig. 2): of all 74 mutated genes identified in coevolved genotypes, only a single gene was found to be mutated in both of these two groups, indicating that their evolutionary trajectories were completely different. Based on these data, we conclude that strong selection resulting from a synergistic coevolution is indeed causal for both the observed divergence in evolutionary trajectories as well as the accelerated accumulation of mutations.

Parallel evolution among replicate lineages is a strong signature of selection favouring the corresponding mutations (*28*). Particularly high levels of parallel evolution were for instance observed in tryptophan-auxotrophic isolates from cocultures: *ompF*, which encodes the outer membrane porin F (OmpF), was found to be mutated in all ten replicate populations. The detected mutations replaced amino acids with longer side chains (i.e. tyrosine, glutamic acid, aspartic acid), which face into the pore-channel, with shorter ones (i.e. alanine, glycine, serine, and cysteine) (see respective SNPs in Supplemental Table 1), thus potentially increasing permeability for exchanged amino acids (*29, 30*). Moreover, 82% of all analysed samples (14 out of 17) carrying mutations in *ompF* were also found to have a mutation in *lrp*. The majority of these mutations caused a frameshift (i.e. ∼57%), thus rendering the leucine-responsive regulatory protein (Lrp) non-functional. Importantly, knockouts of the global regulator Lrp exhibit a selective advantage during stationary phase, which resembles the amino acid starvation experienced by cocultured auxotrophs in our selection experiment. Interestingly, a side-effect of deleting *lrp* is also a reduced ability to transport aromatic amino acids such as tyrosine and tryptophan (*31*), which could explain the strong selection for increased OmpF-permeability in these mutants.

In the case of the cocultured tyrosine auxotrophs, the spectrum of detected mutations was more diverse and thus less similar between strains (Fig. 3B). Nevertheless, also here, striking examples for parallel evolution were observed: In five out of ten populations, a IS5-mediated 10-11 kB deletion was detected within the exact same region comprising 12 genes, amongst them *sspA*. The benefit of lacking the functional stringent starvation protein A (SspA) is particularly interesting, given that a frameshift mutation within *sspA* was additionally detected in another population. In contrast to genotypes with non-functional Lrp, strains lacking SspA exhibit reduced viability during prolonged stationary phase (*32*) by for instance increased sensitivity to acidification (*33*). This detrimental effect needs to be compensated by a yet unknown mechanism. One advantage originating from this mutation is hypermotility due to released H-NS expression (*33, 34*), which potentially increases the likelihood to encounter compatible partners for cross-feeding, when not being part of a multicellular cluster. This effect was likely particularly important, when cocultures experienced low cell densities after being transferred into fresh culture medium during the evolution experiment.

Our results show that mutualistic coevolution increases the rate of molecular change in genomes of synergistically interacting organisms. However, an alternative explanation to account for the observed pattern could simply be demographic differences among experimental groups. Indeed, at the beginning of the evolution experiment, auxotrophic cocultures grew much slower and to a much lower density than populations of the two control treatments (*25*). However, until the end of the evolution experiment, cocultured auxotrophs managed to reduce their generation time as well as to increase their growth rate and the final population density they achieved at the end of a growth cycle. Eventually, coevolved auxotrophs even overhauled auxotrophic monocultures regarding these fitness parameters to finally reach levels that were statistically indistinguishable from the ones of monocultures of prototrophic controls (*25*). Nevertheless, genetic drift in populations of a small size could have led to a genome-wide accumulation of non-adaptive genomic changes, ultimately giving rise to an increased rate of molecular evolution. However, several observations rule out this possibility. First, genotypes of cocultured auxotrophs formed specific branches in the distance tree that were clearly distinct from the ones of control groups (Fig. 2). Second, genomes of coevolved auxotrophs showed the highest level of parallel evolution in comparison to the other experimental groups (Fig. 3A). Third, the vast majority of mutations detected in coevolved auxotrophs were either non-synonymous (clones: 35,8%, populations: 31,2%), non-sense (clones: 9,9%, populations: 10,4%), or insertions as well as deletions (clones: 38,9%, populations: 42,9%) (Supplemental Table 1), suggesting these are adaptive changes. None of these observations would be expected if only demographic differences caused the increased rate of molecular evolution.

A feature that was critical for the evolution of mutualistic cooperation in our experiment was the formation of multicellular clusters that consisted of both types of auxotrophs (*25*). These aggregates most likely enhanced the exchange of amino acids between cells in the otherwise shaken liquid environment (*26, 27*). Under these conditions, the fitness of a given bacterial cell within a cluster is limited by the supply of adaptive mutations that, for example, enhance production levels of the exchanged amino acid. The emergence of such mutations has been shown to cause a positive fitness feedback, in which an increased cooperative investment by one cell is immediately rewarded: enhancing the growth of the corresponding other partner automatically results in an increased supply of the required amino acid in return. Multicellular clusters, in which these mutations arise, will increase in frequency and therefore be transferred to the next growth cycle with a higher probability than clusters lacking these mutations. Thus, competition between clusters that differ in their mutational makeup, can likely explain the enhanced rate of evolution observed in coevolved auxotrophs. This powerful mechanism can therefore account for the increased rate of molecular evolution of mutualistically interacting individuals observed in our study. Interestingly, this situation is strikingly similar to the one mutualistic interactions face in nature. Also here, mutations that enhance growth and reproduction of the whole mutualistic consortia will likely be favoured over consortia lacking these mutations.

Our work identifies coevolution between two mutualistically interacting genotypes as a powerful mechanism to drive molecular change. Strong synergistic benefits resulting from the ecological interaction among individuals set off an evolutionary dynamic, in which strains have to continuously evolve novel adaptive mutations in order to persist against other competing groups. Reciprocal selection pressures emerging from coevolutionary interactions may therefore be causally involved in generating the bewildering diversity in form and function one can observe in mutualistically interacting species.

## Materials and Methods

### Bacterial strains

We used *Escherichia coli* BW25113 (*35*) as the wild type (WT), which was genetically modified by P1 phage transduction (*36*). Derived auxotrophic genotypes contained an in-frame replacement of the targeted amino acid biosynthesis gene (i.e. *trpB* or *tyrA*) with a kanamycin cassette. To allow discrimination of different genotypes on agar plates, the phenotypic marker genes *araDAB* (derived from *E. coli* REL607 (*37*)) and *lacZ* (derived from *E. coli* MG1655 (*38*)) were additionally introduced into WT and auxotrophic strains by P1 transduction. As a result, one set of strains carried the functional alleles for arabinose utilization and β-galactosidase, which appears blue on TA-Agar (*39*) supplemented with 0.1 mM IPTG (Isopropyl β-D-1-thiogalactopyranoside) and 50 µg ml^-1^ X-Gal (5-bromo-4-chloro-3-indolyl-β-D-galactopyranoside), while the other set of WT phenotypes appears red.

### Culture conditions

In all experiments, minimal medium for *Azospirillium brasilense* (MMAB) (*40*) with 0.5 % glucose instead of malate and without biotin was used as culture medium. Cultures were incubated under shaking conditions at 30 °C and 225 rpm. Only monocultures of auxotrophic genotypes were supplemented with tryptophan or tyrosine (150 µM for precultures and 50 µM for the evolution experiment). To start an experiment, bacterial strains were freshly streaked from cryo-stocks on LB agar plates and incubated for 18-24 h at 30 °C. Individual colonies were used as biological replicates to inoculate each 1 ml MMAB of overnight preculture, which were set to an optical density (OD_600nm_) of 2 the next day. Respective aliquots of these cultures were subsequently used to inoculate 4 ml MMAB medium with an initial OD_600nm_ of 0.005. In the case of auxotrophic cocultures, each genotype was inoculated with an initial OD_600nm_ of 0.0025.

### Evolution experiment

Complementary auxotrophic strains (*E. coli ΔtrpB::kan araDAB lacZ,and E. coli ΔtyrA::kan ara-*Δ*lacZ* or the reverse combination of phenotypic labelling) were combined in cocultures to generate a synthetically designed obligate byproduct interaction. To determine effects of genotypic background (prototrophy or auxotrophy, and presence or absence of phenotypic marker genes) on the accumulation of mutations, two control groups contained monocultures of utilized genotypes. Six biological replicates of each generated genotype were used to start the evolution experiment, adding up to twelve monocultures of WT, twelve cocultures of auxotrophs, and 24 monocultures of auxotrophic genotypes (i.e. 12 of each type). Populations were initially transferred every seven days for a total of five transfers, which was followed by 15 transfers every three days, adding up to a total of 80 days or approximately 153 bacterial generations. At the end of each cycle, optical densities were determined in 200 µl culture in microtiter plates by spectrophotometry in a plate reader (Spectramax M5, Applied Biosystems, United States) and 20 µl of culture were transferred into 4 ml of fresh MMAB-medium. Depending on the cycle-length, glycerol stocks (20% glycerol) were prepared each six or seven days and stored at -80 °C. Cocultures were regularly tested for revertant phenotypes that showed prototrophic growth (i.e. were capable to grow on MMAB-Agar without amino acid supplementation). Two out of twelve cocultures were excluded from further analysis due to the evolution of prototrophic phenotypes. Accordingly, also the matching biological replicates in auxotrophic monocultures were excluded from further analysis. In addition, one replicate of *E. coli* Tyr^-^ Ara^-^ Lac^-^ from monocultures was excluded due to contamination. Terminal populations were spread on modified TA agar plates to isolate evolved clones based on colour and colony morphology for whole-genome resequencing. Phenotypic diversity was observed in eight out of ten cocultures, and three out of 19 monocultures of auxotrophs, while all WT monocultures remained phenotypically homogeneous. Isolates were stored at -80 °C until further analysis.

### Genome resequencing and analysis

Evolved populations were sequenced on the level of isolated clones and on the metagenome-level. For this, isolates from terminal populations were incubated in LB medium and whole populations in the respective native minimal medium until the maximum optical density was reached. Genomic DNA was extracted using the Epicentre MasterPure™ Complete DNA & RNA purification kit (MC85200, Biozym Scientific, Germany). Further steps were performed by the Max Planck Genome Centre Cologne, Germany (https://mpgc.mpipz.mpg.de/home/): Quality control of samples was performed on Genomic DNA ScreenTape Analysis® using TapeStation Analysis Software A.02.01 (Agilent Technologies, USA), followed by TruSeq-compatible library preparation. Clonal samples were sequenced on the Illumina HiSeq2500 platform in 100-bp paired-end mode for all WT and coculture samples and in 150-bp paired-end mode for samples from auxotrophic monocultures. CASAVA (v1.8.2) (*41*) and bcl2fastq2 (v.2.18.0.12) (*42*) were used for basecalling and demultiplexing. We used cutadapt (v1.9.1) (*43*) for trimming Illumina sequencing adapters with a minimal overlap of 12. PhiX sequences were identified using bowtie (v1.9.1) (*44*) and BBMap (v37.28) (*45*). We discarded reads with fewer than 15 nucleotides. Only complete read pairs were kept. Observed coverage was approximately 75-fold at quality scores above 30. Sequencing was successfully performed for 65 clonal samples in total. Numbers of sequenced clones depended on the number of observed morphotypes on agar plates (see above). Further analysis of mutations revealed two cases of clones from the same population of cocultures to exhibit identical genome sequences. To avoid pseudoreplication, the corresponding pairs were treated as one. Besides genotypes isolated from derived populations of WT (n = 10), monocultures (n = 22, with Δ*trpB*: n=11 and Δ*tyrA*: n=11), and cocultures of auxotrophs (n = 31, with Δ*trpB*: n = 18 and Δ*tyrA*: n = 13), also the six ancestral genotypes were sequenced to identify mutations that were already present at the beginning of the evolution experiment. Metagenomes were sequenced on the Illumina HiSeq3000 platform using 150-bp single-end mode. To allow quantitative comparison of accumulated mutations without sampling bias, whole populations were analysed from WT populations (n = 6) as well as of populations of auxotrophic monocultures (n = 12, with Δ*trpB*: n=6 and Δ*tyrA*: n=6) and their corresponding cocultures (n = 6). Observed coverage was approximately 1,250-fold at quality scores above 30. Mapping of reads on the published reference genome of *Escherichia coli* BW25113 (CP009273_1) (*46*) and identification of mutations was performed using the BRESEQ-pipeline (*47, 48*). For population samples, the polymorphism mode with the “Polymorphic Read Alignment (RA) Evidence” option “--polymorphism-minimum-coverage-each-strand” was set to 40. Identified mutations and evidence for new junctions (“Unassigned new junction evidence”) were rechecked by verifying individual reads to sufficiently indicate the presence of the mutations and to only map once onto the genome, especially when using the polymorphism mode. If reads indicating a certain mutation were confirmed to map elsewhere in the reference genome with 100% homology by using NCBI nucleotide BLAST (10.1093/nar/gkn201), as in the case of e.g. the highly homologous tRNA-encoding genes, respective mutations were rated as false-positives and excluded from further analysis. Absolute numbers of mutations were determined by counting each mutation regardless of size or structure as a single event. Complex mutations were first resolved in clonal samples as described for BRESEQ (*48*) and further used to successfully resolve all detected complex mutations in population samples by confirming identical architecture (for instance reads to be tiled at identical positions).

One population of auxotrophs that evolved in monoculture carried a non-sense mutation in the gene *mutS*. Inactivation of *mutS* is well-known to cause a hypermutator phenotype that rapidly accumulates increased numbers of single nucleotide polymorphisms (SNPs) (*49*). These *mutS*-induced mutations have mainly neutral or deleterious effects (*50*). Since this study aimed at analysing the effect of synergistic coevolution on the rate of genomic evolution, the corresponding population as well as the respective isolates were excluded from further analysis.

### Quantification of parallel evolution

The Jaccard Index (J) was calculated to estimate parallel evolution at the level of shared mutated genes between samples (*51*). J-values range between zero and one with lower values indicating that fewer mutations occurred simultaneously in the compared samples and larger numbers pointing to an increased similarity between samples. J was calculated for all possible combinations within individually sequenced isolates and within whole population samples, excluding comparisons with the same sample.

### Distance trees

Divergent evolution between experimental groups was analysed with a standard neighbour-joining method and GrapeTree (*52*) was used for visualisation. All identified mutations were summarized in a genotyping matrix that indicated whether a given gene within each sample was either mutated or showed the WT allele. Complex mutations that affected more than one gene were treated as single alleles as well, since these mutations were single evolutionary events that separate a mutant from another lineage.

## Acknowledgments

The authors would like to thank the entire Kost-lab (current and past) for discussion and Wilhelm Boland for support. The Max Planck-Genome-centre Cologne (http://mpgc.mpipz.mpg.de/home/) is gratefully acknowledged for the sequencing of clonal isolates and meta-population samples in this study.

## Author contributions

Conceptualization: DP, CK

Methodology: DP, AH

Investigation: DP

Visualization: DP

Funding acquisition: CK

Project administration: CK

Supervision: CK

Writing – original draft: DP, CK

Writing – review & editing: DP, CK, AH

## Data and materials availability

Sequencing data will be made publicly available by uploading it to a data repository (NCBI) once the article has been accepted for publication.

## Supplementary Materials

Figs. S1 to S2 Tables S1 to S2

