## Supplementary Materials for "Synergistic coevolution accelerates genome evolution"

**This PDF file includes:**

Figs. S1 to S2

**Other Supplementary Materials for this manuscript include the following:**

Tables S1 to S2

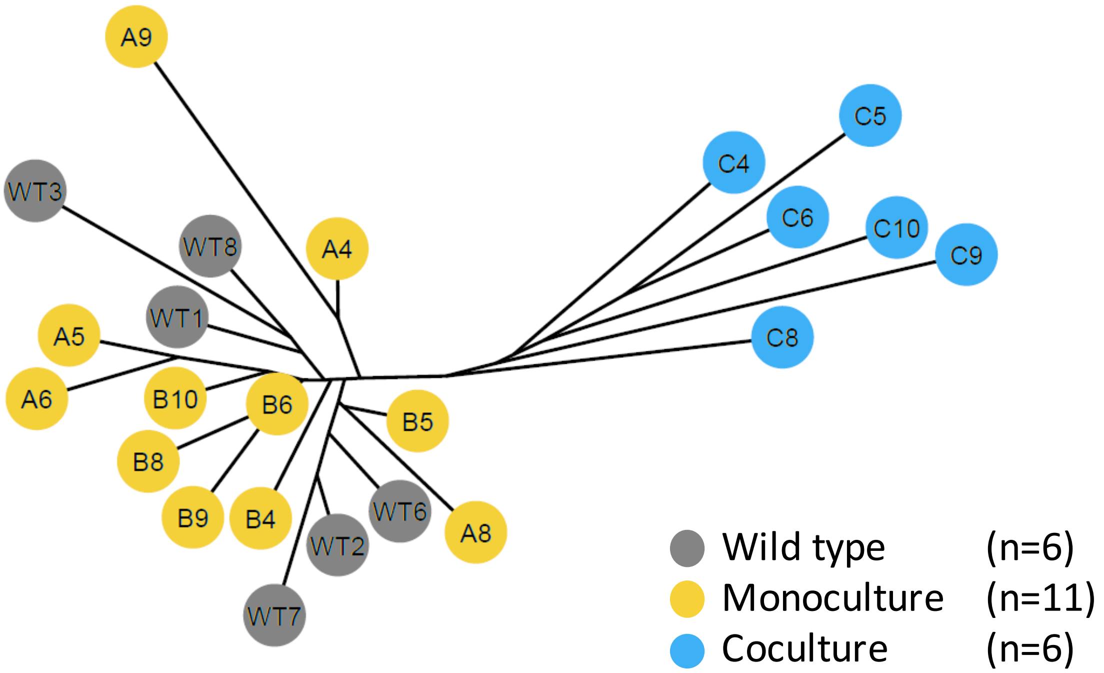


**Fig. S1. Increased divergence of populations of coevolved auxotrophs.** Distance tree of sequenced populations that is based on the neighbour-joining analysis of a population-level genotyping matrix. Nodes are colour-coded by experimental group. Evolutionary trajectories differ between control groups (WT = wildtype, M = monocultures of auxotrophs, C = cocultures of auxotrophs). Labels containing similar numbers indicate a common ancestor among auxotrophic populations.

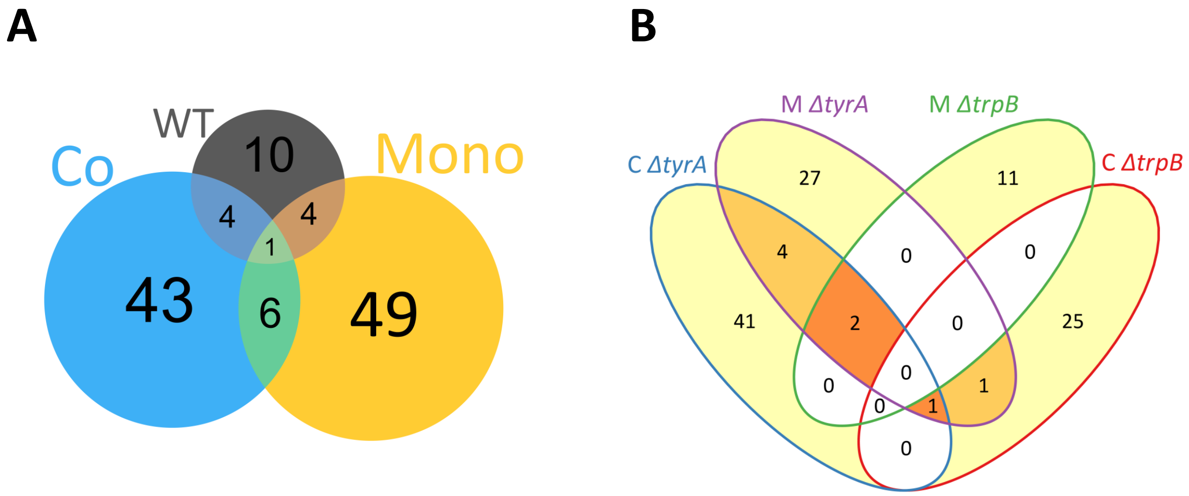


**Fig. S2. Experimental groups share few mutated genes.** Numbers in Venn-diagrams represent total counts of different genes carrying a mutation or having been deleted. (**A**) Shared and unique mutated genes in populations of wild type (WT), monocultures of auxotrophs (Mono), and cocultures of auxotrophs (Co). (**B**) Shared and unique mutated genes in auxotrophic clones isolated from cocultures (C Δ*tyrA*, C Δ*trpB*) and monocultures (M Δ*tyrA*, M Δ*trpB*).

**Supplementary Tables (not included in this PDF)**

**Table S1. Overview over all mutations that have been identified in experimental groups on the level of isolated strains.**

**Table S2. Overview over all mutations that have been identified in experimental groups on the level of whole population samples.**
